# Spatio Temporal Attentional EEGNet: An Enhanced Deep Learning Model for Cognitive Workload Detection

**DOI:** 10.1101/2025.09.26.678835

**Authors:** Sayan Ghosh, N. Nikhil Siddharth, J. Gowthami, Dheemanth Chakravathi, N. Maheedhar Sai

## Abstract

Deep learning has emerged as a powerful tool for extracting meaningful patterns from electroencephalography (EEG) signals, particularly for mental workload recognition in cognitive neuroscience. This study proposes STA-EEGNet (Spatio Temporal Attentional EEGNet), an enhanced variant of the EEGNet architecture that integrates spatio temporal attention mechanisms to improve EEG signal decoding. The model is trained and evaluated on the STEW: Simultaneous Task EEG Workload Dataset, demonstrating superior classification accuracy compared to existing approaches. Heatmap analyses of key convolutional filters in TSA-EEGNet network during testing reveal distinct spatiotemporal activation patterns associated with varying workload levels. High workload is characterized by increased activation in discriminative filters and reduced or inverted responses in suppressive filters, primarily over frontocentral and parietal regions. These findings highlight TSA-EEGNet’s ability not only to achieve state-of-the-art performance but also to provide interpretable insights into the neural dynamics underlying cognitive workload.

## I. Introduction

EEG records the brain’s electrical activity by capturing signals produced through neuronal oscillations, offering a direct measure of brain function [1]. It enables the real-time tracking of neural markers linked to mental effort, including variations across frequency bands [2, 3–4]. Studies have consistently demonstrated that distinct EEG patterns are associated with different workload levels, highlighting the effectiveness of EEG for continuous monitoring of cognitive states [5–6]. EEG-based mental workload classification approaches traditionally rely on feature extraction from time, frequency, and time–frequency domains[7,9–10]. Commonly used features include spectral power in theta (4–8 Hz), alpha (8–12 Hz), and beta (13–30 Hz) bands, event-related potentials (ERPs), and nonlinear measures such as entropy and fractal dimension, which are then fed into classifiers like Support Vector Machines (SVM), Linear Discriminant Analysis (LDA), and Random Forests for state discrimination [9–10]. For example, Berka et al. [7] demonstrated that theta power increases and alpha power decreases are robust indicators of elevated workload, enabling accurate classification in aviation and monitoring tasks. Recent advancements have increasingly focused on deep learning models capable of learning discriminative spatiotemporal representations directly from raw EEG signals, thereby minimizing reliance on hand-crafted features [3–4, 11]. Convolutional Neural Networks (CNNs) have been extensively employed to extract spatial patterns across EEG channels [8], while Recurrent Neural Networks (RNNs), particularly Long Short-Term Memory (LSTM) units, are utilized to model temporal dependencies [11]. Bashivan et al. [8] proposed a recurrent-convolutional approach by transforming EEG into multi-spectral topographic images, achieving significant performance gains in workload classification tasks. Likewise, BiLSTM-LSTM framework has been applied to improving both classification accuracy and model interpretability in cognitive load assessment [11].

The aim of utilizing EEG signals for cognitive load detection is to assess a person’s mental workload in real time by analyzing brain activity during task performance. This method is particularly valuable in healthcare, where it can help monitor cognitive load in individuals with neurological conditions such as dementia, traumatic brain injury, and psychosis, etc. The structure of this paper is organized as follows: Section II. A provides a detailed description of the data acquisition process and the preprocessing techniques used. Section III describes the architecture of the classifier network. SectionS IV and IV.A present the results of the network’s performance evaluation, along with a comparative analysis and discussion on its biological resemblance. Finally, Section V concludes the work by summarizing the proposed approach and highlighting potential directions for future research.

## II. Method

### A. Data Acquisition and Preprocessing

The STEW (Simultaneous Task EEG Workload) database [12] comprises EEG recordings from 48 participants, each completing two experimental conditions: the ‘No’ task and the SIMKAP (Simultaneous Capacity) task [13]. In the ‘No’ task, participants remained in a comfortable seated position with their eyes open and performed no specific activity. In the SIMKAP task, participants completed a multitasking and stress assessment developed by Schuhfried GmbH, designed to evaluate cognitive capacity under simultaneous task demands. Each task was recorded for three minutes of EEG data, and to minimize transitional effects, the first and last 15 seconds of each recording were discarded. Following each task, participants rated their mental workload (MWL) on a scale from 1 to 9. EEG signals were recorded using the Emotiv headset with a sampling frequency of 128 Hz and a 16-bit A/D resolution. The recordings included 14 electrodes (AF3, F7, F3, FC5, T7, P7, O1, O2, P8, T8, FC6, F4, F8, AF4) placed according to the 10–20 international system. The classification problem can be approached in two ways: 1. Three-class scenario – workload is categorized as poor (scaling 1–3), moderate (scaling 4–6), or high (scaling 7–9). 2. Two-class scenario – workload is grouped into normal (scaling 4–6) and high (scaling 7–9). The scaling was done based on the subject feedback form. In this study, we focus on the two and three-class classification problem. The EEG data underwent band-pass filtering between 0.1 and 40 Hz, followed by Independent Component Analysis (ICA) for artifact removal, and subsequent z score normalization. The continuous 2.5-minute recordings were segmented into 1-second windows (128 samples) with 50% overlap. Although the dataset originally included 48 participants, three were excluded due to missing ratings. Of the remaining participants, data from 36 (80%) were used for training and 9 (20%) for testing.

## III. Model Architecture

We propose a Temporal-Spatial Attention EEGNet model for EEG-based mental workload classification (fig 1). The input EEG signals first pass through a temporal convolution layer (TC) to extract time-domain features, followed by depthwise and separable convolution layers (DC and SC) to capture channel-specific and combined spatial-temporal patterns. A spatial attention module (SAM) then highlights the most informative electrode regions. The attended features are processed through a TC2 layer (Conv1D layer) for final feature refinement. The model outputs classification results indicating mental workload levels. The mathematical description of STA-EEGNet has been discussed in the supplementary file but the novel finding of the spatial attentional map has been elaborated in Section III. A.

**Fig. 1.**
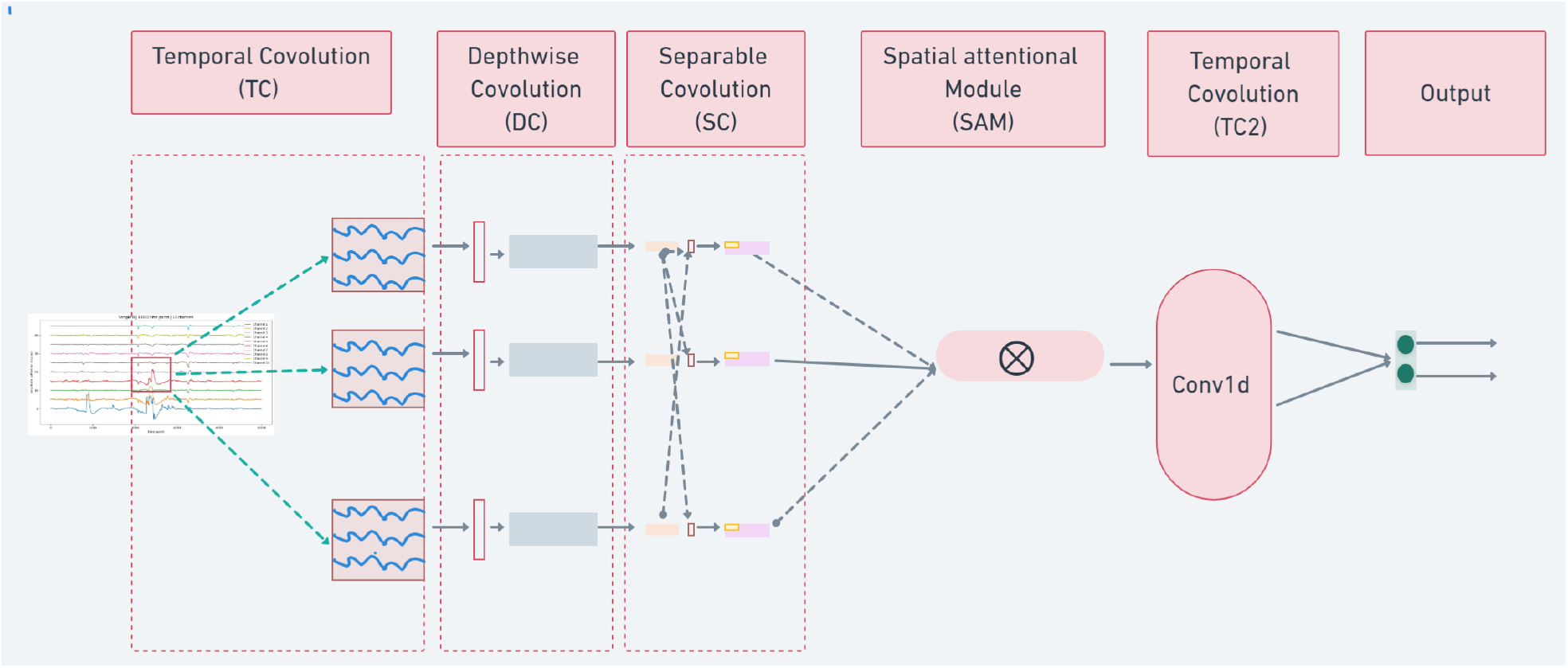
Architecture for Spatio-Temporal Attentional EEGNet.

### A. Spatial Attention Module (SAM)

Mean over channel (or filters) is described in eqn. (1).

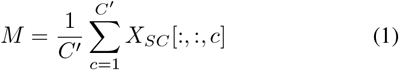

*Convolution over mean of filter:*

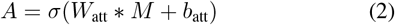

Where, *A* ∈ ℝ^*H×W ×*1^. Here, *C*^*′*^ denotes the number of feature channels in *X*_*SC*_, where *SC* refers to the separable convolution layer. *X*_*SC*_ represents the activation output of the SC layer. A 2D convolution is then applied M (Mean over channel), where *W*_att_ and *b*_att_ denote the learnable convolution weights and bias term, respectively. And *σ* is the sigmoid activation function. Finally, element-wise multiplication (eqn. (3)) is performed to obtain the attention map (*X*_10_).

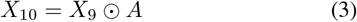

## IV. Result

From a total of 45 subjects, 80% of the data (36 subjects) was used for training and the remaining 20% (9 subjects) for testing. The evaluation was performed in a cross-subject manner, ensuring that data from the same subjects did not appear in both training and testing sets. Each subject’s recording was 2.5 minutes long and segmented into 1-second chunks, resulting in 13,455 total samples. Of these, 10,764 segments from 36 subjects were used for training and 2,691 segments from 9 subjects for testing. Codes are available on GitHub, as mentioned in the supplementary file.

To assess model robustness, the experiment was repeated 20 times, and the results were averaged over multiple runs. The proposed model achieved an average classification accuracy of 92.37% for the 2-class problem and 82.31% for the 3-class problem (Table II), surpassing state-of-the-art performance.

**TABLE I.** Network parameter.

|  |  |
| --- | --- |
| **Input** | Raw EEG signal, Batch, 14, 128, 1; channel=14, time points=128 |
| **Temporal Convolution (TC Layer)** | Kernel Length=64, Filters=8, Shape: batch, 14, 128, 8 |
| **Depth Convolution (DC Layer)** | Size of depth-wise filter: 14,1; depth multiplier=2, activation: Elu, average pooling: (1,4) |
| **Separable convolution (SC Layer)** | Filters: 16, size of filter: (1, 16), padding='same', activation: Elu, average pooling: (1,8) |
| **Spatial Attention module (SAM Layer)** | Number of filter: 1, size of filter: (1,1); activation: sigmoid; padding='same'; kernel regularizer=l2(1e-4) |
| **Conv1D layer (TC2 Layer)** | Filter: 64, kernel size: 3, activation: ReLU |
| **Output Layer** | 2/3, activation: softmax (Two-class or 3-class problem) |
| **Learning Rate** | 0.0001 |
| **Optimizer** | adam |
| **Loss function** | Categorical cross-entropy |
| **Dropout rate** | 0.2 |
| **Epoch** | 300 |
| **Trainable parameter** | 4,932 |
| **Batch size** | 32 |

**TABLE II.** Comparison Table for STEW Dataset.

| Study | Accuracy |
| --- | --- |
| [11] BiLSTM, LSTM with PSD, AR coeff., approx. entropy, Hurst exponent first-order statistics | 86.33 (2-class), 82.57 (3-class) |
| [12] STEW Dataset, PSD feature | 3-class classification: 69% |
| [15] EEGNET | 84.33±1.03 (2-class); 83.64±1.33 (3-class) |
| [14] OVG, DT, SVM | 89.6 (2-class), 79.5 (3-class) |
| Proposed Network (STA EEGNet) | 2-class: 92.37%, 3-class: 82.31 |

### A. Neurological Resembles

The heatmap analysis (fig 2) reveals distinct spatiotemporal EEG patterns associated with different cognitive workloads across several key convolutional filters. In Filter 0, Low Workload elicits strong activation in channels AF3, F7, F3, FC5, T7 during 10–40 samples, whereas High Workload shows weaker responses shifted to 80–100 samples. Filter 3 demonstrates marked workload sensitivity, with Low Workload producing positive peaks in channels AF3, F7, F3, FC5 and FC6, F4, F8 around 20–40 samples, while High Workload generates stronger, sustained activation in the same channels during both 20–40 and 90–110 samples. Filter 5 exhibits more negative activation under High Workload, particularly in channels AF3, F7, F3, FC5 and P8, T8 between 20–50 samples, suggesting suppression of these brain regions when cognitive load is high. Filter 7 mirrors Filter 3’s pattern, with High Workload amplifying peaks in channels AF3, F7, F3, FC5 and P8, T8, especially in the 20–40 sample window. These effects predominantly involve fronto-central and parietal electrodes, regions linked to attention and working memory, with early windows (20–40 samples) likely reflecting stimulus-locked responses and later windows (90–110 samples) indicating sustained or secondary processing. Overall, High Workload tends to enhance activation in discriminative filters (3 and 7) while reducing or inverting activity in suppressive filters (e.g., 5).

**Fig. 2.**
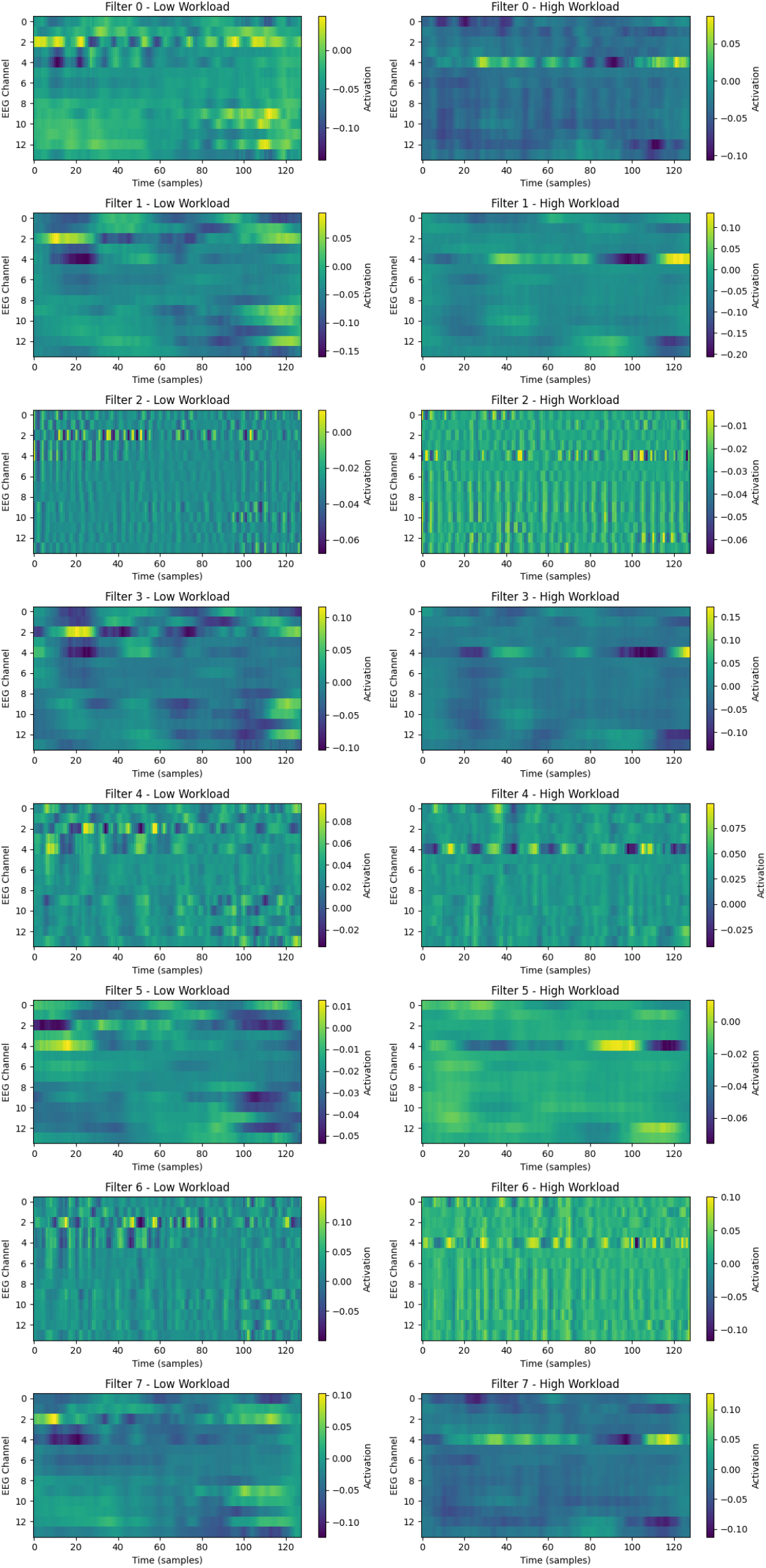
Activation maps for filters 0–7 under low and high workload conditions.

## V. Conclusion and future direction

The proposed STA-EEGNet demonstrated superior performance in cognitive workload classification, achieving 92.37% accuracy for the two-class problem on the STEW dataset, outperforming conventional EEGNet[15] and other state-of-the-art methods (Table II). Heatmap analyses revealed distinct spatiotemporal patterns between low and high workload conditions, with fronto-central and parietal electrodes playing a dominant role. Early time windows (~ 20–40 samples) likely reflected stimulus-locked processes, while later windows (~ 90–110 samples) captured sustained or secondary cognitive processing. Discriminative filters (3 and 7) exhibited amplified activation under high workload, whereas suppressive filters (e.g., 5) showed reduced or inverted activity, underscoring the model’s ability to capture both excitatory and inhibitory neural dynamics relevant to workload differentiation.

Future work can extend this approach by integrating frequency-domain attention mechanisms to capture oscillatory features explicitly (e.g., alpha, beta rhythms), applying explainable AI techniques such as layer-wise relevance propagation for deeper interpretability, and validating across larger, more diverse EEG datasets to ensure generalizability. Additionally, combining STA-EEGNet with real-time adaptive BCI systems could enable closed-loop workload monitoring in high-stakes environments such as aviation, driving, and neurorehabilitation. Longitudinal studies could further explore how workload-related EEG patterns evolve with training, fatigue, or neurological conditions, opening new avenues for personalized cognitive state tracking.

## Acknowledgment

We sincerely appreciate SRM AP for supporting this work with computational resources.

